# Isolation and characterization of three lytic podo-bacteriophages with two receptor recognition modules against multidrug-resistant *Klebsiella pneumoniae*

**DOI:** 10.1101/2024.05.19.594906

**Authors:** Liang Huang, Xueting Huang, Tongfei Zhao, Jingren Zhang, Ye Xiang

## Abstract

The multidrug-resistant nosocomial pathogen *Klebsiella pneumoniae* is considered as one of the major threats to public health. Recent studies showed that bacteriophages could be used as an alternative to antibiotics for treating infections caused by multidrug-resistant *K. pneumoniae.* Here we isolated and characterized three lytic bacteriophages Kp7, Kp9 and Kp11 of *K. pneumoniae*. Transmission electron microscopy analysis showed that all three phages have an isometric head of approximately 60 nm in diameter and a short noncontractile tail. Host specificity, one-step growth curve, tolerance to pH and temperature changes were characterized for Kp7, Kp9, and Kp11. Mass-spectrum and bioinformatical analysis identified two different types of tail fibers in each phage. The tail fiber proteins gp52 of Kp7, gp43 of Kp9 and gp46 of Kp11 are homologous and have robust enzymatic activity in digesting purified serotype K2 capsule of *K. pneumoniae*. However, tail fiber proteins gp51 of Kp7, gp42 of Kp9 and gp45 of Kp11 have diverse sequences and show weak or no enzyme activity in digesting purified serotype K2 capsule. Genetic knockout and biochemical assays indicated that the capsule is essential for the infection of Kp7, Kp9 and Kp11. Spot tests showed that 15 clinical isolates of drug-resistant *K. pneumoniae* with the serotype K2 capsule can all be infected by Kp7, Kp9 and Kp11 but with different susceptibility. Further mechanistic studies showed that transcriptional inhibition at the early stage of phage infection determines the susceptibility of different *K. pneumoniae* isolates.

## 1. Introduction

The long-term usage and especially the abuse of antibiotics promote a positive selection of multidrug-resistant bacterial pathogens, which pose a great challenge to public health and raise an urgent requirement for the development of new generation of antibiotics. The Gram-negative *Klebsiella pneumoniae* is non-motile and rod-shaped, and a common nosocomial pathogen responsible for a large number of infections every year^1^. *K. pneumoniae* belongs to the Enterobacteriaceae family and as an opportunistic bacterial pathogen is a natural inhabitant of upper respiratory and intestinal tract. Most of *K. pneumoniae* strains are coated with thick polysaccharide capsules. The *K. pneumoniae* capsules have highly diverse structures and compositions, and have more than 79 serotypes^2^. The capsule is closely related to the virulence of *K. pneumoniae* and strains with serotype K1 or K2 capsule are the major cause of liver abscesses^3–5^. Antibiotic resistance of *K. pneumoniae* is largely due to its ability of frequently acquiring antimicrobial resistance (AMR) genes, through either horizontal gene transfer (HGT) or de-novo mutations under selection pressure^6^. Recent *K. pneumoniae* isolates are shown to resist almost all the antibiotics including carbapenem, which is usually the last line antibiotic used for treating bacterial infections^7–9^.

Bacteriophages, also known as phages, are potent viruses that infect and lysate specific bacterial hosts. Due to the rapid emerging of multidrug-resistant bacteria, antibiotics are predicted to be exhausted in the near future. Phage therapies regained increasing attention in recent years^10,11^. Phage therapies are highly specific and efficient. However, most bacteriophages infect and lysate only several or even one strain of a specific bacterium. Thus, the stringent specificity of bacteriophages is also considered as the major shortage of phage therapy. Phage cocktails and bacteriophage encoded endolysins or depolymerases were developed to broaden the antibacterial spectrum of phages^12–15^. Alternatively, precise engineering of phage receptor binding proteins (RBPs) has been successful in several phages to broaden the antibacterial spectrum^16,17^. More RBPs and bacteriophages with multiple types of RBPs need to be characterized and molecular mechanisms that govern the assembly and function of the RBPs need to be illustrated, for precise engineering and practical application of phages^18,19^. Bacteria develop resistance to phage infection through different mechanisms, which also hinder therapeutical applications of phages^20^.

In this study, we isolated three phages Kp7, Kp9, and Kp11 that target *K. pneumoniae.* Genome sequencing and bioinformatical analysis of Kp7, Kp9, and Kp11 showed that all of them have a dsDNA genome with short direct terminal repeats, which is encapsidated in an icosahedral capsid attached to a short non-contractile tail. Two different types of tail fibers were identified in each phage and show different enzyme activities in digesting the serotype K2 capsule. The stability test showed that Kp7, Kp9, and Kp11 are relatively stable at pHs ranging from 4 to 11 and at temperatures ranging from 37 to 60 □. Genetic assays proved the essential role of capsules for phage infection. Fifteen clinical *K. pneumoniae* isolates with the serotype K2 capsule showed different susceptibility to Kp7, Kp9, or Kp11 infection. Further investigation identified a mechanism of transcriptional inhibition for phage resistance in insensitive *K. pneumoniae* strains.

## 2. Materials and Methods

### 2.1 Bacterial strains

*K. pneumoniae* strain ATCC 43816 (TH13157 in this study) from the American type culture collection (ATCC) was used for initial phage isolation and purification. The clinical *K. pneumoniae* isolates and the genetically modified *K. pneumoniae* strains (TH13863, TH14082, TH15628) were from the collections of Dr. Jingren Zhang.

### 2.2. Phage isolation and purification

Water samples (10 mL each) collected at different locations in Beijing were centrifuged under a G force of 4000 ×g for 10 minutes at 4 □. The supernatant of each collection was filtered through a 0.22 μm filter and then was concentrated with an Amicon Ultra-4 30 kDa cutoff filter. Initial screenings of bacteriophages were performed by using classical double-layer agar overlay technique^21^. Briefly, 100 μL fresh *K. pneumoniae* cell culture (OD_600_=0.6, 5×10^8^ CFU/mL) was mixed with 100 μL of the concentrated water sample. The mixture was added to 4 mL 0.7 % soft LB agar and was then overlaid onto the surface of an LB agar plate. After the solidification of the soft agar, the plate was inverted and cultured at 37 □ for 12 hrs.

To amplify the phage from a single plaque, phages in the plaque were picked up with a pipette tip and then transferred into LB culture of *K. pneumoniae* (OD_600_=0.6, 5×10^8^ CFU/mL). The infected cells were growing in a 37 □ orbit shaker at a speed of 220 rpm and were lysed by phages in approximately 2 hours after infection. The lysate was centrifugated at 4000 ×g for 15 mins and then filtered with a 0.22 μm filter to remove cell debris. The filtered supernatant was used as a stock for large scale phage amplification. The purification procedure was carried out by using the isopycnic CsCl gradient ultracentrifugation^22^. Briefly, the filtered supernatant was centrifuged at 4 □ and a speed of 25000 rpm for 2 hrs with a Beckman SW50.2Ti rotor. The pellet was resuspended in TMS buffer (50 mM Tris-HCl, pH 7.8; 100 mM NaCl and 10 mM MgCl_2_). The resuspended phages were further purified by using an 80 % (w/v) isopycnic CsCl gradient with a SW55Ti rotor at 25000 rpm and 4 □ for 12 hrs. After the centrifugation, the opaque band of phage was taken out with a syringe. The purified phages were exchanged into the TMS buffer to remove the CsCl and stored in TMS buffer at 4 □. The titer was determined by using plaque assays.

### 2.3. Electron microscopy

The concentrated phage at a concentration of 10^11^ PFU/mL was negatively stained and analyzed with an FEI Tecnai Spirit transmission microscope. Briefly, an aliquot of 4 μL purified phage was applied onto a glow-discharged carbon film supporting grid for 1 min, then the liquid was removed with a filter paper and washed twice with 7 μL 2 % uranyl acetate each time. The grid was stained for 1 min with 7 μL 2% uranyl acetate after the wash and then air-dried for 3 mins. The grids were checked on an FEI Tecnai Spirit TEM operating at 120 kV with a magnification of 52,000×.

### 2.4. Genomic DNA extraction and restriction enzyme digestion

The genome DNA of Kp7, Kp9, and Kp11 was extracted and purified from the concentrated phages (10^12^ PFU/mL). In brief, 20 μL of the concentrated phage was mixed with 500 μL TE buffer (50 mM Tris pH 7.8, 10 mM EDTA) and proteinase K, and the mixture was incubated at 70 □ for 1 h. Then the solution was added with 500 μL 25:24:1 phenol/chloroform/isopropanol and thoroughly mixed by gentle inversion. Next, the solution was centrifuged at 12,000 ×g for 5 min and approximately 250 μl of the upper layer solution was carefully transferred to a new tube. After adding 750 μL prechilled 100 % ethanol, the genomic DNA was pelleted by centrifugation and rinsed twice with 75 % ethanol. Finally, the genomic DNA was resuspended with 50 μL ddH_2_O (∼160 ng/μL) and sent for sequencing and agarose gel analysis. To verify the genome sequencing results, the purified genomic DNAs were analyzed by restriction endonuclease digestions (*Bsiw* I for Kp7, *Age* I for Kp9, and *BspQ* I for Kp11) (NEB Cat. R0553V, R0552V and R0712S). Each reaction contained 500 ng of the purified genomic DNA and the corresponding enzyme and was incubated for 1 hour at 55 □ for *Bsiw* I, 50 □ for *BspQ* I, and 37 □ for *Age* I. The digested products were analyzed by electrophoresis with 0.7 % agarose gels.

### 2.5. Genome sequencing and bioinformatic analysis

The purified phage genomic DNA was sequenced by using the whole genome de novo sequencing method. To determine the exact termini of the genomes, raw sequencing reads were analyzed by using the software PhageTerm^23^. The genome sequences were assembled and annotated by using RAST (http://rast.nmpdr.org/)^24^. Open reading frames (ORFs) were predicted with the software GeneMarkS^25^. Conserved protein domains were identified by using the CD-Search tool at NCBI^26^. The regulatory elements such as tRNA, bacteriophage-specific promoters and rho-independent transcriptional terminators were identified by using the software tRNAScan^27^ and PhagePromoter^28^, respectively. For comparative genomic analysis, the assembled genomes were analyzed by Blastn and tBlastx against non-redundant protein sequence (nr) database and sequence similarities were detected by using the software Easyfig^29^.

### 2.6. Measurements of pH and temperature stability

To invest the pH and thermal stability of the isolated phages, pHs ranging from 3 to 11 and three temperatures 37 □, 50 □, and 60 □ were chosen. For pH stability assays, the phages were diluted in 1 mL TMS buffer at pHs ranging from 3 to 11 to a final concentration of 10^8^ PFU/mL. After an incubation at 25 □ for 60 minutes, the titer was determined by using plaque assays. For the thermal stability assays, phages at a concentration of 10^8^ PFU/mL were incubated at different temperatures for 20 mins, 40 mins and 60 mins, respectively. Then, the phage titer was determined accordingly. All the assays were repeated in triplicate.

### 2.7. One-step growth curve

To determine the bactericidal ability of the isolated phages, one-step growth curves were measured for Kp7, Kp9, and Kp11. Briefly, the refreshed *K. pneumoniae* strain TH13157 was grown to the exponential phase (OD_600_=0.6, 5×10^8^ CFU/mL). To adjust the MOI to 0.01, the cells were infected by mixing 1 mL bacteria with 10 μL purified phage (5×10^8^ PFU/mL). After an adsorption time of 3 mins, the mixture was centrifugated at 13,000 ×g for 2 mins and the supernatant was abandoned to get rid of unabsorbed phages. The pellet was resuspended with 1 mL fresh LB medium. Then, aliquots (10 μL each) of bacterial culture were taken every 5 mins to 10 mins for phage titer determination by using double-layered plaque assays until 90 minutes post infection. All the assays were repeated in triplicate. The burst size is defined as the ratio between the number of phages produced during the plateau period to the number of infected bacteria at the latent period, which is equal to the ratio between the phage titer in the plateau period and the phage titer in the latent period.

### 2.8. SDS-PAGE gel and mass spectrum analysis

The purified phage virions with the SDS-PAGE gel loading buffer were boiled for 20 mins at 100 □. After centrifugation, the samples were analyzed by gradient SDS-PAGE (4-20 %) gels. Protein bands were visualized by silver staining. For identification of possible viral structural proteins, each visible band was cut off from the gel and sent for mass spectrum analysis.

### 2.9. Phylogenetic tree analysis

To build the phylogenetic trees, sequences of tail fiber protein were subjected for homologous protein search by using Blastp with an E-value cutoff of 1×10^3^. Then, multiple sequence alignments were conducted by using ClustalX and the phylogenetic tree was built based on the neighbor-joining method with the software MEGA7^30,31^.

### 2.10. Purification of tail fiber proteins

Tail fiber protein genes, that encode gp51 (1-668 aa) and gp52 (1-575 aa) of Kp7, gp42 (1-777 aa) and gp43 (1-524 aa) of Kp9, and gp45 (1-404 aa) and gp46 (1-524 aa) of Kp11, were PCR amplified from phage genome DNA and were cloned into the vector pETDuet1 via sites *BamH* I and *Hind* III that introduce an N-terminal 6×His tag to the recombinant proteins. The plasmids were transformed into the strain *E. coli* Rosetta (DE3) for recombinant protein expression. Expression of the recombinant proteins was induced with 1 mM isopropyl β-D-1-thiogalactopyranoside (IPTG) at 16 □ for 22 hrs. The recombinant proteins produced were affinity purified using cobalt-charged BD TALON resins and eluted with a buffer containing 20 mM HEPES at pH 7.5, 150 mM NaCl, 5 mM MgCl_2_ and 500 mM imidazole. The eluted proteins were concentrated and further purified via a Superdex 200 column (GE) running in a buffer containing 20 mM HEPES at pH 7.5, 150 mM NaCl and 5 mM MgCl_2_. The peak fractions were collected and used for enzyme activity assays.

### 2.11. Purification of the *K. pneumoniae* K2 capsule

The K2 capsule of *K. pneumoniae* was extracted by using a modified method adapted from previous publications^32^. Briefly, *K. pneumoniae* (TH13157, TH13044, TH13091, and TH13148) cells recovered on LB agar plates containing 3 % sheep blood were seeded in 50 mL LB and cultured overnight at 37 □ with shaking (60 rpm). Then the cultures were cooled in ice bath and harvested by centrifugation at 17,700 ×g, 4 □ for 30 mins. After removing the supernatant, the cell pellets were resuspended in 5 mL buffer containing 0.1 % zwittergent 3-14 and 50 mM citrate at pH 4.5. The resuspension was incubated in a 42 □-water bath for 30 mins and then was centrifuged at 17,700 ×g, 4 □ for 5 mins. The supernatant containing the capsule was concentrated via an Amicon Ultra-4 100 kDa filter and further purified via a Superdex 6 column (GE) running in a buffer containing 20 mM HEPES at pH 7.5, 150 mM NaCl and 5 mM MgCl_2_. The peak fractions containing the capsules were collected and concentrated for testing the enzyme activity of tail fiber proteins.

### 2.12. Enzyme activities of the tail fiber proteins

The enzyme activity of the tail fiber proteins was tested by incubating 10 μL of the purified capsules with 5 μL each of the serial dilutions of the purified tail fiber proteins at 37 □ for 1 hour. The reactions were quenched by adding the SDS loading buffer and boiled at 96 □ for 10 mins. Then the samples were subjected for SDS-PAGE gel analysis. The capsules were visualized by Alcian blue staining, as described previously^33^. Briefly, the gels were rinsed in water containing 25 % ethanol and 10 % acetic acid for 30 mins. Then the gels were stained in dark for 15 mins at 50 °C in a staining solution containing 0.125 % Alcian blue, 25 % ethanol and 10 % acetic acid. The gels were distained overnight at room temperature with 25 % ethanol and 10 % acetic acid.

### 2.13. Host range determination

Forty-two drug-resistant clinical isolates of *K. pneumoniae* used in this study are listed in Table S1. Fifteen *K. pneumoniae* strains with the serotype K2 capsule used in this study are listed in Table S2. Prior to host range determination, *K. pneumoniae* strains were refreshed by plate streaking. For the spot assays^34^, 100 μL fresh cell culture of *K. pneumoniae* (OD_600_=0.6-0.8) was mixed with 3 mL 0.7 % soft LB agar. After the solidification of the agar, 1 μL phage stock (10^10^ PFU/mL) was spotted onto the plate. Susceptibility of *K. pneumoniae* strains or isolates to phage infections was determined by using spot assays with serial dilutions of phage stocks, of which the concentrations are in the range from 10^6^ PFU/mL to 10^10^ PFU/mL. To investigate the function of capsules in phage infection, genetically modified *K. pneumoniae* strains, which include TH13863, TH14082 and TH15628, were used.

### 2.14. Constructing capsule deficient mutants of *K. pneumoniae*

*K. pneumoniae* strain TH13863 with the unmarked deletion of capsular polysaccharide (CPS) biosynthesis gene cluster was constructed with the strain TH13157 by using a CRISPR-based mutagenesis approach as described^35^. The 25-kb CPS region contains 18 genes required for the synthesis of CPS, from the 5’ to the 3’ including *galF, cpsACP, wzi, wza, wzb, wzc, wcuF, wcuD, wclX, wzy, wzx, wcsU, wcaJ, wcaJ, gnd, manC, manB* and *ugd*. Full deletion of the CPS region in TH13157 was accomplished by three sequential steps. Step1: the 5’ end of the CPS region from *galF* to *wzc* was deleted to construct the strain TH13692 as follows. The plasmid pCasKP was electroporated into TH13157 to generate strain TH13179, which harbors the Cas9 nickase and was used as the recipient strain. The sgRNA expression plasmid pTH13416 with the spacer Pr15206/Pr15207 that targets the gene *galF* was constructed. The donor repair template was constructed by fusion PCR that fuses a 1000 bp upstream sequence of *galF* (amplified with Pr15230/Pr15578) and a 1000 bp downstream sequence of *wzc* (amplified with Pr15579/Pr15580). Then, the sgRNA expression plasmid and the repair template were co-transformed into the recipient strain TH13179 to generate mutant TH13692, which has *galF* to *wzc* fragment deleted and was used for deleting the middle part of the CPS region. Step2: the middle part of the CPS region from *wcuF* to *wcaJ* was deleted in TH13692 to construct TH13811. The sgRNA expression plasmid pTH13609 with the spacer Pr15425/Pr15426 that targets *wcuF* and the corresponding repair template were co-electroporated into TH13692 to generate TH13811 that harbors the deletion of the *galF* to *wcaJ* fragment. The repair template was generated by fusion PCR that fuses the 1000 bp upstream sequence of *galF* (amplified with Pr15230/Pr15578) and the 1000 bp downstream sequence of *wcaJ* (amplified with Pr15579/Pr15580). Step3: the 3’ end of CPS region from *gnd* to *ugd* was deleted in TH13811 to generate ΔCPS mutant TH13863. The sgRNA expression plasmid pTH13847 with the spacer Pr15592/Pr15593 that targets *gnd* and the repair template that contains the fused 1000 bp upstream fragment of *galF* (amplified with Pr15230/Pr15231) and the 1000 bp downstream fragment of *ugd* (amplified with Pr15232/Pr15233), were co-electroporated into TH13811 to generate ΔCPS mutant TH13863. All primers used to construct the ΔCPS mutant TH13863 were listed in Table S3.

### 2.15. Mechanisms of resistance to phage infection

To explore the intrinsic mechanism involved in the susceptibilities of different isolates to phage infection, phage infection assays were performed for sensitive and insensitive strains. Briefly, the refreshed *K. pneumoniae* strain was grown to the exponential phase (OD_6000_=0.6, 5x10^8^ CFU/mL). To adjust the MOI to 1, four parallel tubes were set with 1mL bacteria and 10 μL purified phage (5x10^10^ PFU/mL) in each tube. After an adsorption time of 3 mins, the mixture was centrifugated at 13,000 ×g for 2 mins and the supernatant was abandoned to get rid of unabsorbed phages. The pellet was resuspended with 1 mL fresh LB medium. Then, one tube of bacterial culture was centrifugated at 13,000 ×g for 2 mins to collect the infected cells. The supernatant was abandoned while the pellets were quickly frozen by liquid nitrogen (The time point was set 0 min post phage infection). Tubes of bacterial culture were harvested every 5 mins until 15 mins post infection. All collected cell pellets were quickly frozen by liquid nitrogen and stored at -80 □. All the assays were repeated in triplicate.

The total bacterial and phage RNA was extracted by using the GeneJET™ RNA Purification Kit (Thermo). Briefly, the cell pellets were pretreated with TE buffer containing 0.4 mg/mL lysozyme. Then the total RNA was extracted and reversed transcribed into cDNA for quantitative PCR by using the PrimeScript RT Reagent Kit (TaKaRa). Quantitative real-time reverse transcriptase PCR (qRT-PCR) was performed using the iTaq Universal SYBR^@^ Green Supermix (Bio-Rad). The following primer pairs were used to amplify the target genes: Pr15170/Pr15171 (bacterial gene *16s rDNA*), Pr15172/Pr15173 (phage gene *orc*), Pr15174/Pr15175 (phage gene *RNA polymerase*). The *16s rDNA* gene was used as a reference for normalization of phage related genes expression levels at different time points post phage infection. The primer sequences were listed in Table S4. All the assays were repeated in triplicate.

## 3. Results

### 3.1. Phage isolation and morphological features

We collected water samples at different locations in Beijing and the collected water samples were used for screening phages against the ATCC 43816 *K. pneumoniae* strain by using double-layer plaque assays (Figure 1A). We observed plaques on the plates incubated with water samples collected from the Sand River, South Sand river of Beijing, and the river near the Tsinghua University hospital. Phages from three plaques surrounded by obvious opaque halos on the plates were then picked with toothpicks and were amplified with *K. pneumoniae* growing in liquid medium. The amplified phages are named Kp7, Kp9, and Kp11, respectively. All these phages were purified by using isopycnic CsCl gradient centrifugation and were examined with an FEI Tecnai Spirit transmission electron microscope (TEM). The results showed that all the three isolated bacteriophages have an icosahedral head and a short non-contractile tail, indicating that they all belong to the family Podoviridae (Figure 1B). By measuring the negatively stained phage particles, the diameters of the icosahedral heads are approximately 65 nm, 61 nm and 62 nm for Kp7, Kp9 and Kp11, respectively.

**Figure 1.**
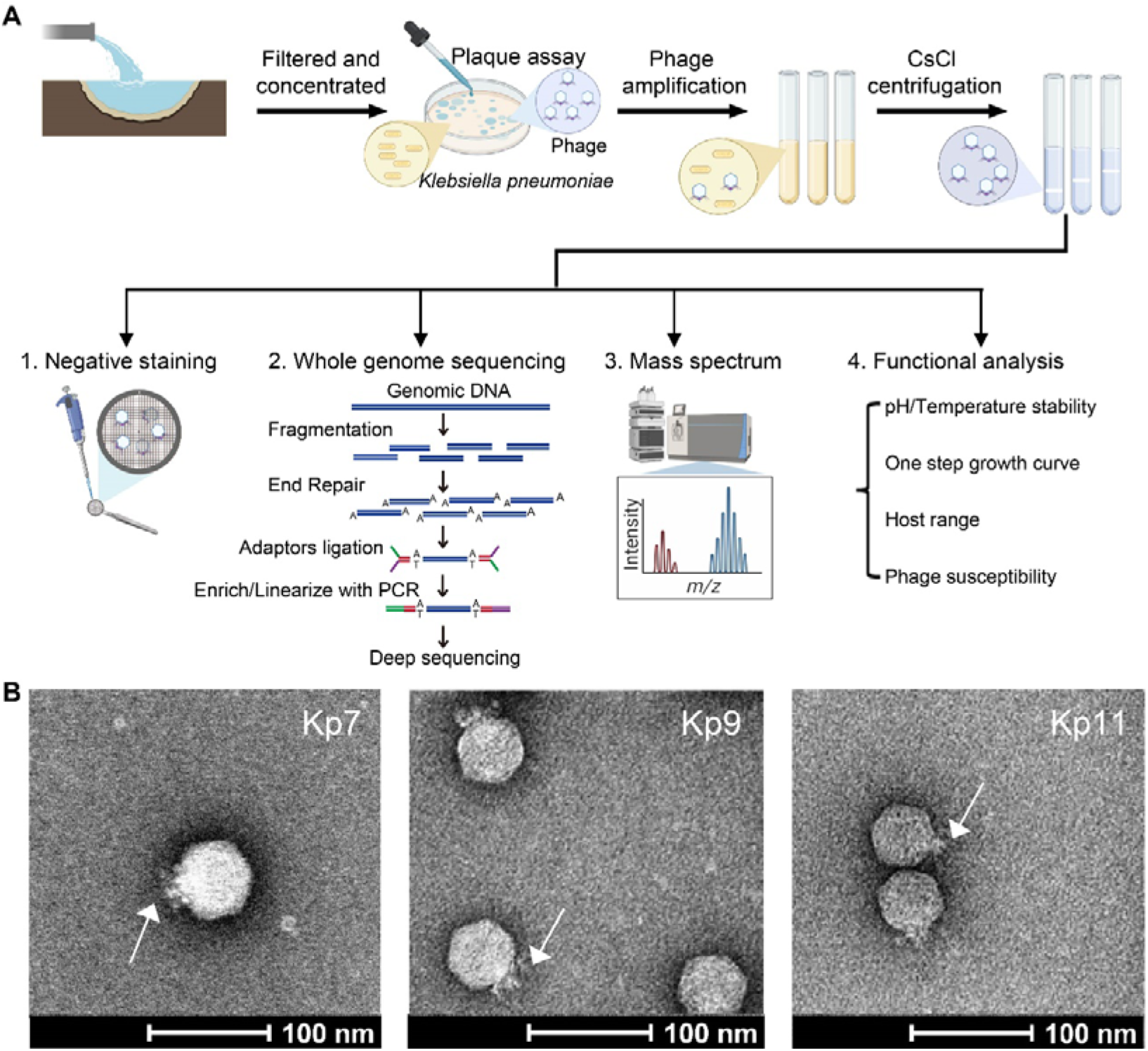
Phage isolation, identification, and characterization. (A) A schematic diagram showing the workflow for the isolation, identification, and characterization of *Klebsiella pneumoniae* phages. (B) Transmission electron micrographs showing negatively stained particles of Kp7, Kp9, and Kp11. The scale bar represents 100 nm. The white arrows indicate the short tails of the isolated phages.

### 3.2. Genomic size and whole genome sequencing

To identify the isolated phages, genomes of Kp7, Kp9 and Kp11 were extracted and sent for sequencing. The assembled genomes were featured with repeating sequences located at both the 5’ and 3’ termini of the genome. These repeating sequences are known as direct terminal repeats, which are probably signals for genome packaging initiation and termination^36^ and are common in T7-like bacteriophage genomes. The terminal repeats analyzed and assembled by using the software PhageTerm^23^ showed that the length of the repeating sequences is 238 bps, 177 bps, and 179 bps for Kp7, Kp9 and Kp11, respectively (Figure 2A).

**Figure 2.**
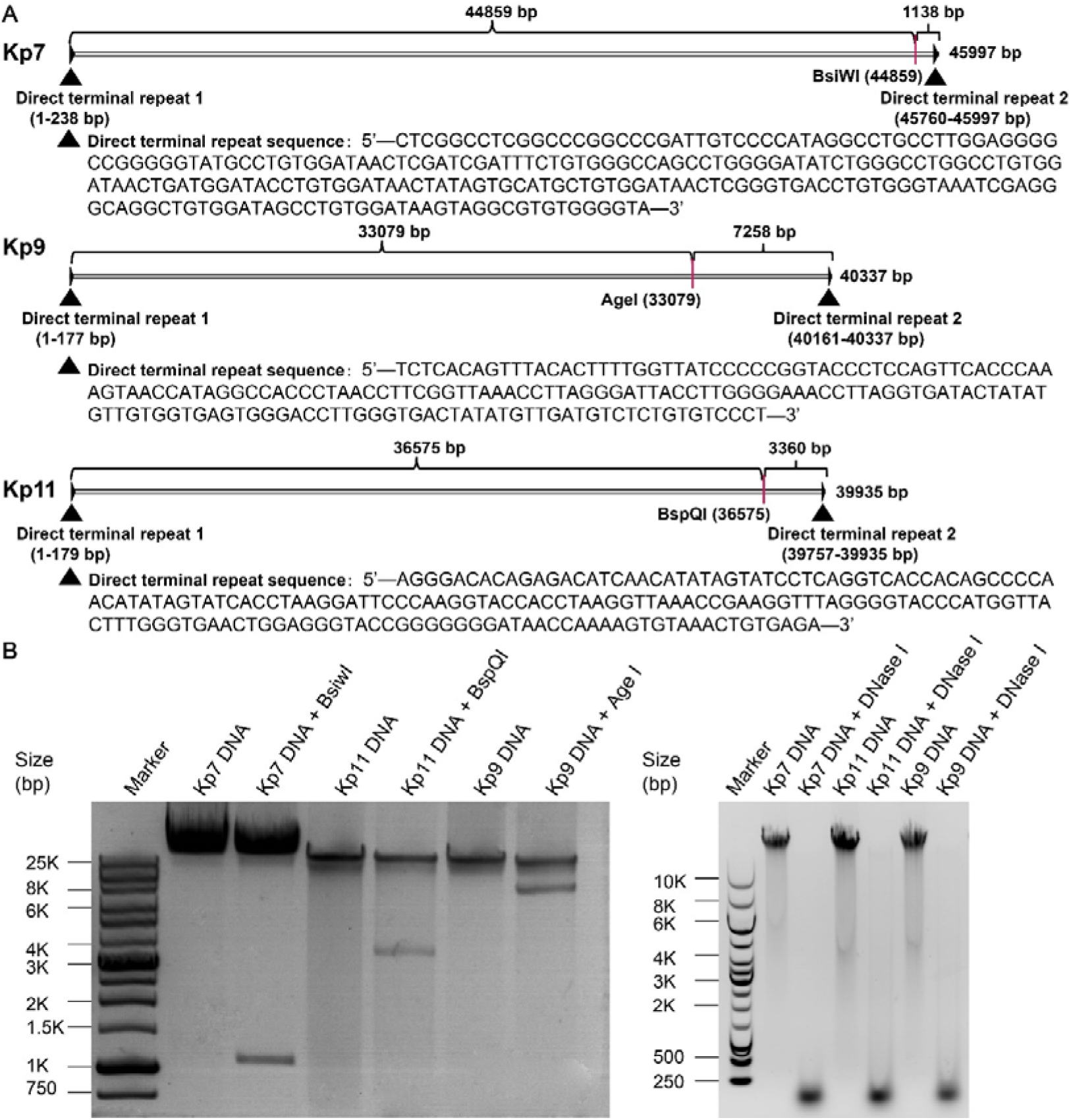
Genome sequencing and validation. (A) Diagrams showing direct terminal repeats identified^23^ in the genomes of Kp7, Kp9, and Kp11. The restriction endonuclease sites indicated were used for validating the sequencing results. The black triangles indicate the positions of the direct terminal repeats, of which the sequences are listed below the diagrams. (B) Enzyme digestions of the genomic DNAs showing fragments with the expected size and features of linear dsDNA.

The assembly results show that Kp7 has a linear dsDNA genome of 45997 bps with a G+C content of 51.9%, Kp9 has a linear dsDNA genome of 40337 bps with a G+C content of 53.1%, and Kp11 has a linear dsDNA genome of 39935 bps with a G+C content of 53.1% (Table S5). Annotation of the coding regions performed by using the RAST annotation server^24^ showed that Kp7, Kp9 and Kp11 have 53, 47 and 50 predicted open reading frames (ORFs), respectively. Further bioinformatic analysis indicates that Kp9 and Kp11 share significant similarities to the reported *K. pneumoniae* phage KP32 that was isolated with the ESBL(+) *K. pneumoniae* strains^37^, whereas Kp7 has only weak similarities with the reported *K. pneumoniae* phages. The sequencing results were further validated by using selected restriction enzymes and DNase I, and the results showed expected cleavage patterns of linear dsDNA genomes (Figure 2A and 2B).

### 3.3. Thermal and pH stability and one-step growth curve of Kp7, Kp9 and Kp11

The pH stability assays showed that Kp7, Kp9, and Kp11 all have the highest titer at pH8. Kp7, Kp9, and Kp11 can form plaques after being treated with buffers in the pH range of 4-11 (Figure 3A). However, the titers of all the phages dropped to zero at pH3 (Figure 3A). The temperature stability assays indicated that the titers of Kp7, Kp9, and Kp11 keep relatively stable at 37 □ but dropped for approximately tenfold after a short time incubation at 60 □ (Figure 3B).

**Figure 3.**
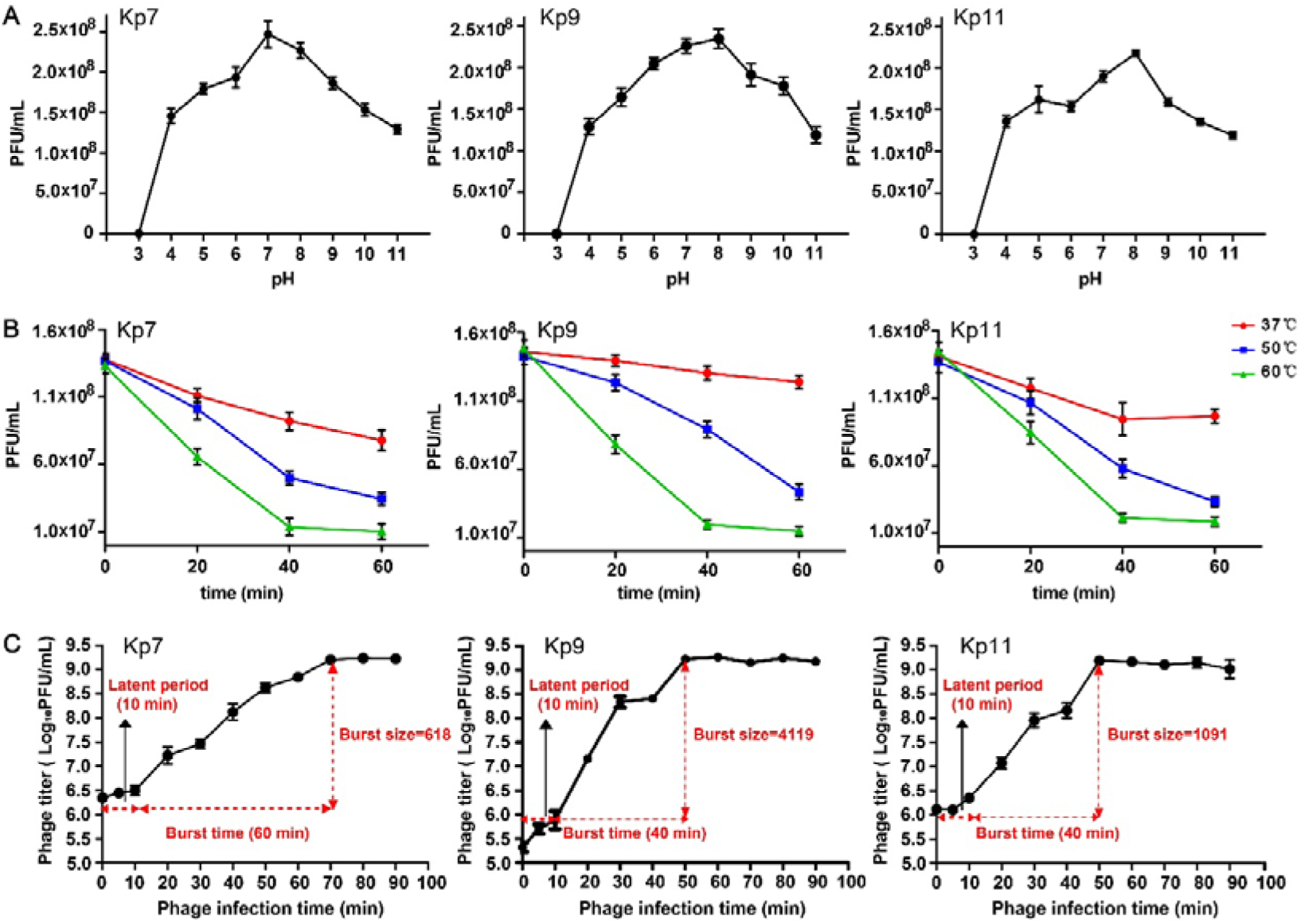
Thermal stability, pH stability, and one-step growth curves of Kp7, Kp9, and Kp11. (A) pH stability of Kp7, Kp9, and Kp11. (B) Thermostability of Kp7, Kp9, and Kp11. (C) One-step growth curves of Kp7, Kp9, and Kp11. The burst size is ratio between the phage titer at the plateau period and the phage titer at the latent period. Values in all the assays are the mean values of three measurements.

The measured one-step growth curves of these phages showed that Kp7, Kp9, and Kp11 have a short and similar latent period of approximately 10 minutes but with different burst sizes (Figure 3C). The burst time of Kp7 is longer (∼ 60 minutes) than these of Kp9 and Kp11 (∼ 40 minutes). The burst sizes of Kp7, Kp9, and Kp11 calculated are 618, 4119, 1091 phage particles per infected cell, respectively (Figure 3C).

### 3.4. Annotation and identification of structural and nonstructural proteins

To identify the structural components of Kp7, Kp9, and Kp11, purified phages were subjected for SDS-PAGE gel and mass spectrum analysis. The SDS-PAGE gels were silver stained. The protein patterns of Kp7 on the SDS-PAGE gel are different to these of Kp9 and Kp11 (Figure 4A). The major band of Kp7 with a molecular weight of ∼40 kDa is identified as the HK97-like major capsid protein gp35. Other bands are identified as the internal virion proteins (gp38, gp39, and gp40), connector (gp33), minor capsid proteins (gp32, gp45, gp46, and gp47), and tail proteins (gp36, gp37, gp41, gp51, and gp52), respectively, through mass spectral analysis and blast search (Figure 4A). Nonstructural proteins are annotated by bioinformatic analysis (Figure 4B). Similarly, structural and nonstructural proteins of Kp9 and Kp11 are also identified and annotated (Figure 4A). Compared to Kp7, both Kp9 and Kp11 lack the minor capsid proteins and the tail adaptor protein, but have two additional structural proteins (gp30 and gp31 for Kp9, gp33 and gp34 for Kp11), of which the functions are predicted to be associated with the correct assembling of the head or the tail^38^. Of note, two different types of tail fiber proteins were identified for all three phages (Figure 4).

**Figure 4.**
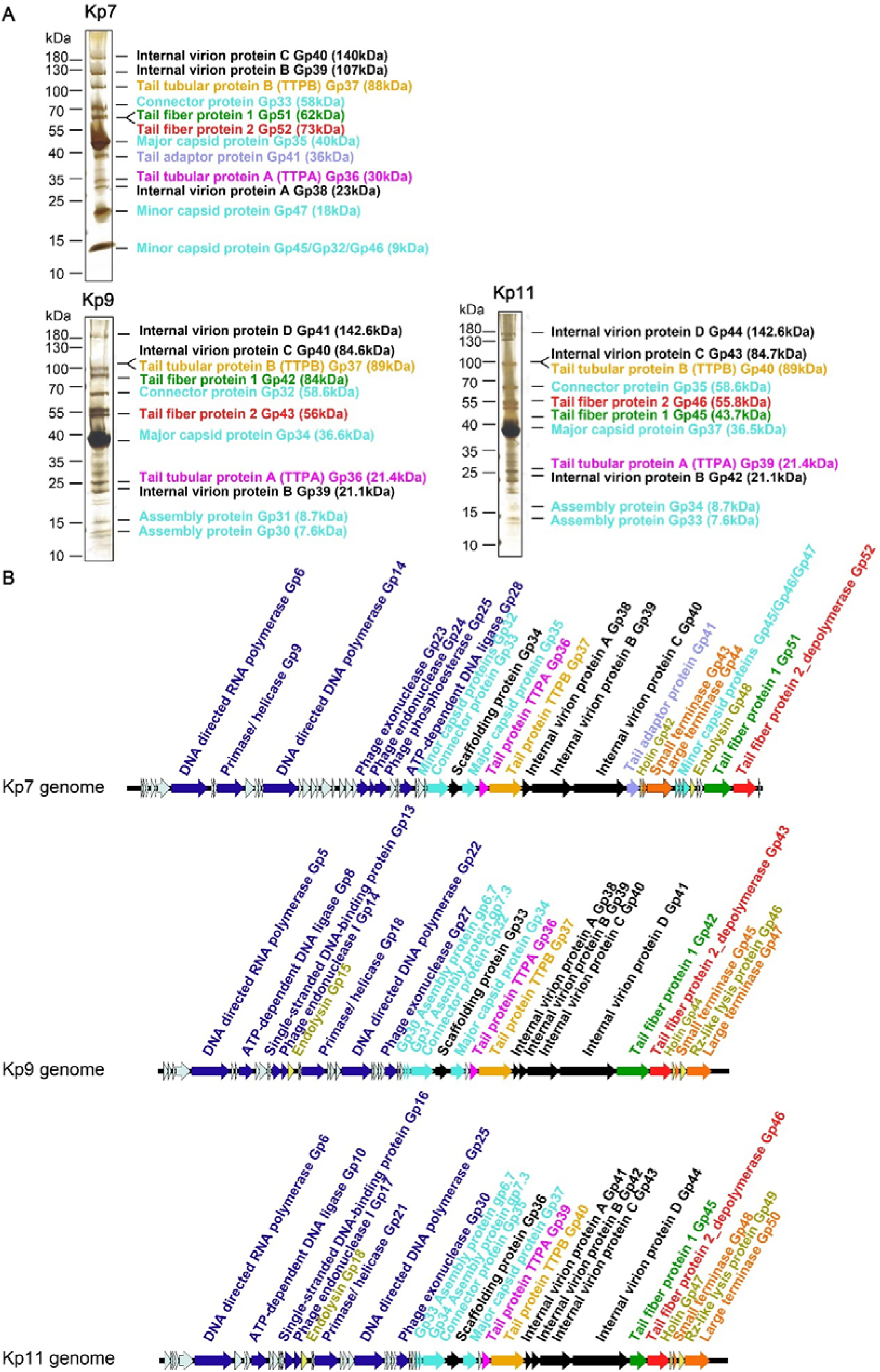
Structural protein identification and ORF annotation of Kp7, Kp9, and Kp11. (A) SDS-PAGE gel analysis of the isolated phage virions. The gels were silver stained. Visible bands on the gels were cut separately for mass spectrum analysis and the results are listed on the right of the corresponding bands. (B) Diagrams showing the linear genome maps of Kp7, Kp9, and Kp11. Arrows represent the predicted ORFs and the direction of arrow represents direction of translation. Different colors represent possible different functions of the predicted ORFs in the life cycles of the phages. ORFs colored in blue are predicted to be involved in phage replication. ORFs colored in black and cyan are predicted to be involved in phage capsid assembly. ORFs colored in magenta, gold, purple, green and red are predicted to be involved in assembling phage tails.

### 3.5. Determination of the host tropisms

To investigate the host specificity of the isolated phages, spot tests were performed with the purified phages and 42 drug-resistant clinical *K. pneumoniae* isolates that have diverse capsules of different serotypes. The results showed that Kp7, Kp9 and Kp11 have a narrow host tropism and infect only two strains among the 42 clinical isolates (Table S1). Both the two susceptible isolates have the serotype K2 capsule as that of the ATCC 43816 *K. pneumoniae* strain used for screening (Table S1). Furthermore, fifteen *K. pneumoniae* isolates with the serotype K2 capsule were used for spot tests with Kp7, Kp9, and Kp11. The results showed that all the fifteen *K. pneumoniae* isolates are susceptible to Kp7, Kp9, and Kp11 infection (Table S2). A series of dilutions of phage stock were then used to test the susceptibility of these K2 capsule isolates to phage infection and the results showed that the fifteen *K. pneumoniae* isolates have quite different susceptibility to Kp7, Kp9 and Kp11. However, sequence analysis of the locus that encodes the capsule synthesis related enzymes showed only minor differences among these strains, suggesting that factors other than the capsule determine the susceptibility to phage infections.

To further identify the roles of the capsule in Kp7, Kp9, and Kp11 infection, phage susceptibility assays were performed with a genetically modified *K. pneumoniae* strain TH13863, of which the entire capsule locus was knocked out from the standard ATCC 43816 *K. pneumoniae* strain (TH13157). The results showed that neither Kp7, Kp9 or Kp11 can infect TH13863 as expected (Table 1 and Supplementary Figure S1). However, phage infections were resumed by using the genetically modified strain TH15628, in which the K2 capsule locus of a non-relevant strain TH12887 was complemented into the TH13863 strain. Of note, phage infections can not be resumed by using the genetically modified strain TH14082, in which the K1 capsule locus of a non-relevant strain TH12908 was complemented into the TH13863 strain (Table 1 and Supplementary Figure S1). Interestingly, the susceptibility of TH15628 to the phages is similar to that of its scaffold TH13157 but not its capsule donor TH12887, further indicating that although the capsule is determinant for phage infection, other factors rather than the capsule determine the susceptibility to phage infection.

**Table 1.**
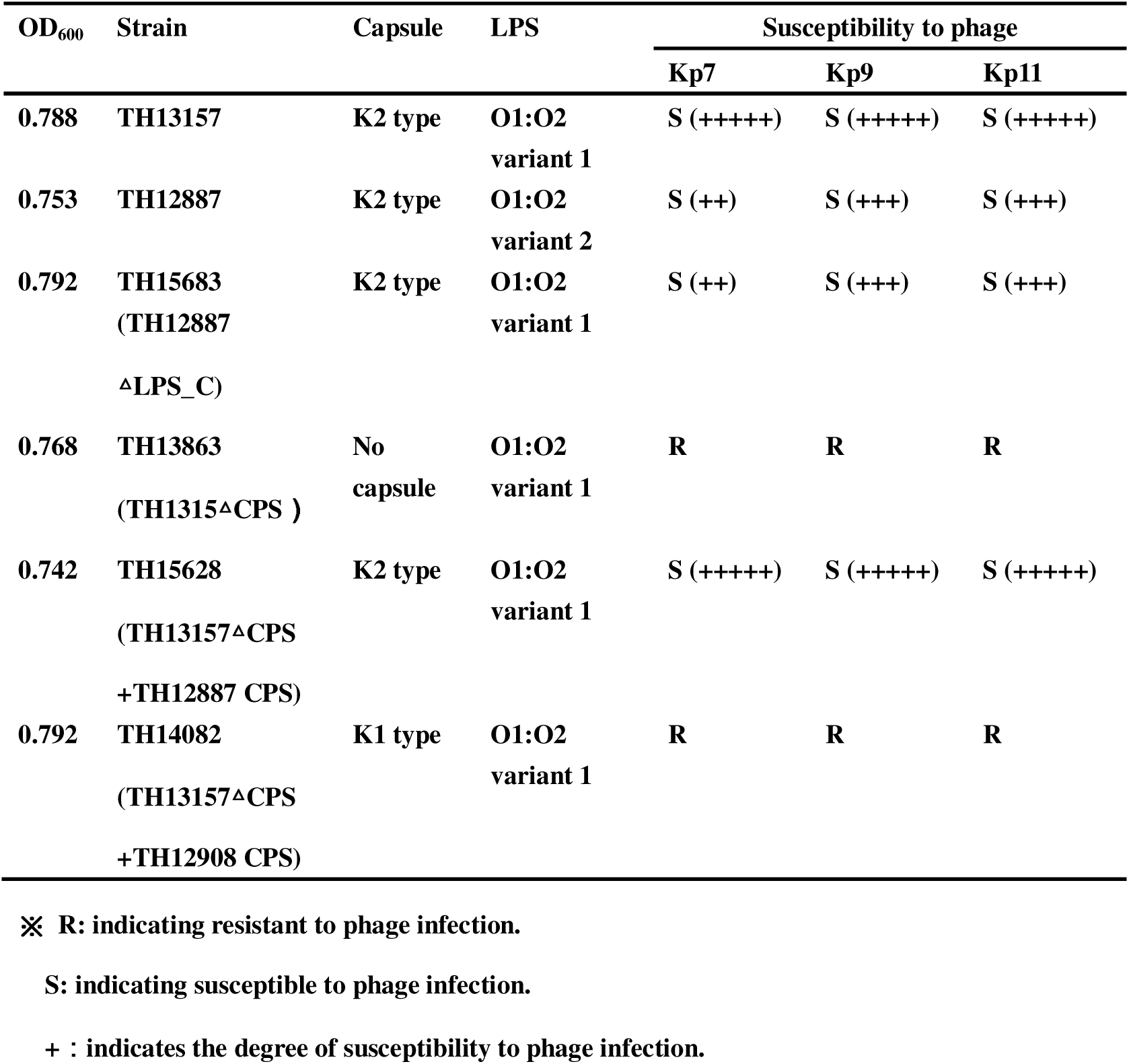
The determinant role of capsule in phage infection.

| OD<sub>600</sub> | Strain | Capsule | LPS | Susceptibility to phage |  |  |
| --- | --- | --- | --- | --- | --- | --- |
|  |  |  |  | Kp7 | Kp9 | Kp11 |
| 0.788 | TH13157 | K2 type | O1:O2<br>variant 1 | S (+++++) | S (+++++) | S (+++++) |
| 0.753 | TH12887 | K2 type | O1:O2<br>variant 2 | S (++) | S (+++) | S (+++) |
| 0.792 | TH15683<br>(TH12887<br>ΔLPS_C) | K2 type | O1:O2<br>variant 1 | S (++) | S (+++) | S (+++) |
| 0.768 | TH13863<br>(TH1315<sup>Δ</sup>CPS ) | No<br>capsule | O1:O2<br>variant 1 | R | R | R |
| 0.742 | TH15628<br>(TH13157<sup>Δ</sup>CPS<br>+TH12887 CPS) | K2 type | O1:O2<br>variant 1 | S (+++++) | S (+++++) | S (+++++) |
| 0.792 | TH14082<br>(TH13157<sup>Δ</sup>CPS<br>+TH12908 CPS) | K1 type | O1:O2<br>variant 1 | R | R | R |

✖ **R: indicating resistant to phage infection.**

### 3.6. Comparisons of the tail fiber proteins

We identified two different types of tail fiber proteins in each of the three isolated phages, suggesting that Kp7, Kp9, and Kp11 may use the two different types of tail fibers for infecting different strains or may need the synergistic function of the tail fibers for infection. Sequence comparisons showed that tail fiber proteins gp52 of Kp7, gp43 of Kp9, and gp46 of Kp11 are similar to each other (Supplementary Figure S2) and to that of the *Escherichia* phage CBA120 tail spike protein 4 (orf213), which has a pyramid like structure and can degrade the O-antigen of *Escherichia coli* O78^39^. A constructed phylogenetic tree showed the close relationship between gp43 of Kp9 and gp46 of Kp11 (Figure 5A). However, tail fiber proteins gp51 of Kp7, gp42 of Kp9 and gp45 of Kp11 have completely different sequences comparing to that of the pyramid-like tail fibers. A constructed phylogenetic tree showed that gp51 of Kp7 is far from other known tail fibers of the *K. pneumoniae* phages (Figure 5B), whereas gp42 of Kp9 and gp45 of Kp11 are similar to each other and share certain sequence homology to that of the tail fiber protein gp37 from *K. pneumoniae* phage KP32 which can degrade the K3 serotype capsule^40^ (Supplementary Figure S3).

**Figure 5.**
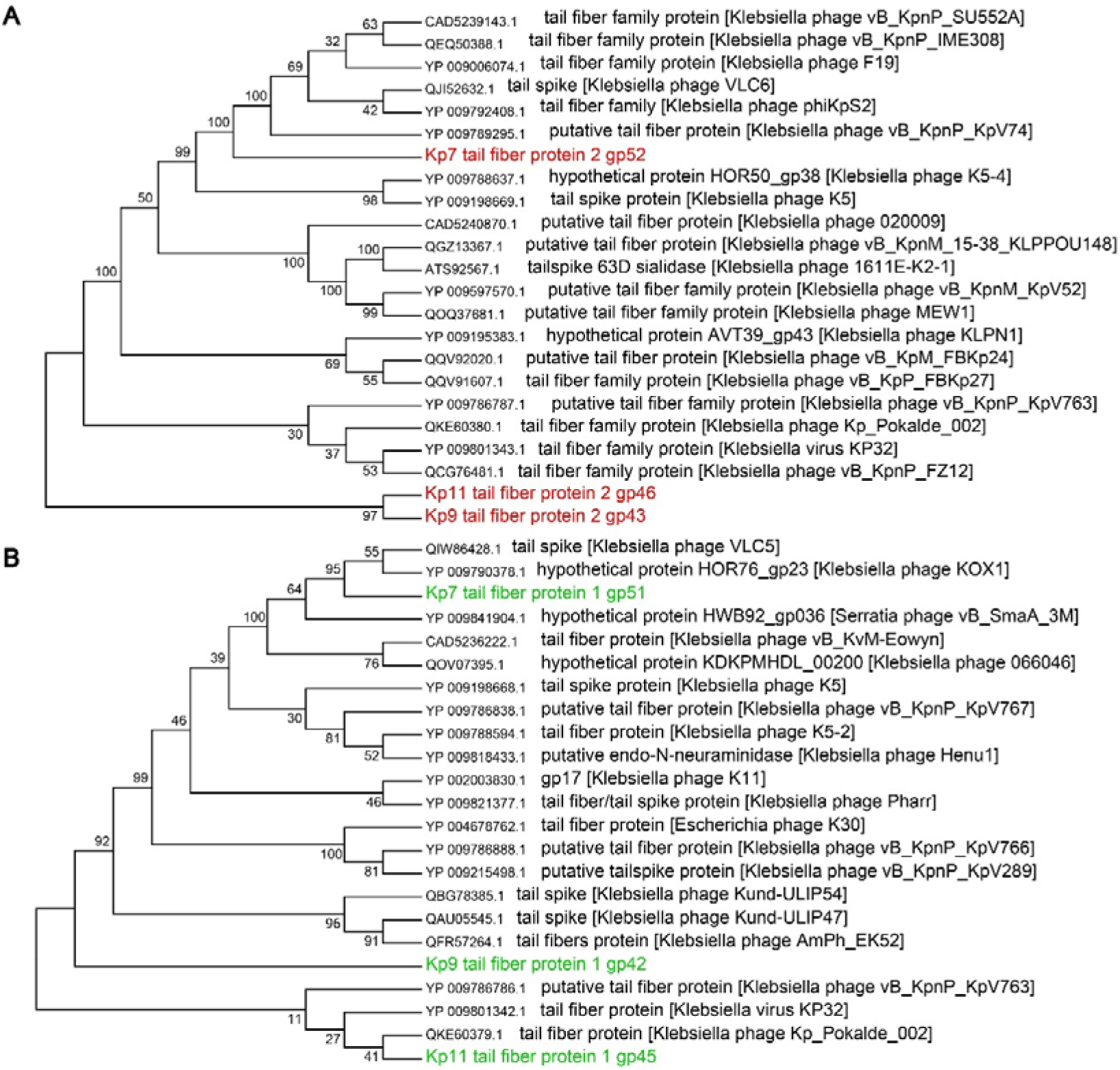
Phylogenetic trees of Kp7 tail fiber proteins gp51 and gp52. Phylogenetic neighbor-joining trees of Kp7 tail fiber gp52 (A) and tail fiber gp51 (B). Tail fiber proteins were blasted for their homologs and sequence alignments were performed by using ClustalX. The software MEGA7 was utilized for constructing the phylogenetic trees.

To understand the function of different tail fiber proteins during phage infection, tail fiber proteins gp51 and gp52 of Kp7, gp42 and gp43 of Kp9, gp45 and gp46 of Kp11 were produced and purified (Figure 6A). SDS-PAGE gel analysis with Alcian blue staining showed that gp52 of Kp7, gp43 of Kp9, and gp46 of Kp11 have robust enzyme activity in degrading the purified K2 capsule, whereas gp51 of Kp7 showed weak enzyme activity, and gp42 of Kp9 and gp45 of Kp11 showed no enzyme activity against the purified K2 capsule (Figure 6B-D). Given the low sequence homology between gp51 of Kp7 and other tail fiber proteins and its weak enzyme activity against the K2 capsule, gp51 may be a newly undefined enzyme that recognizes yet to be identified polysaccharide in the capsules. Besides, plaque assays indicated that the recombinant gp52 instead of the recombinant gp51 of Kp7 could effectively block the formation of the opaque haloes around plaques and completely block the phage infection when supplemented at a higher concentration (Supplementary Figure S4).

**Figure 6.**
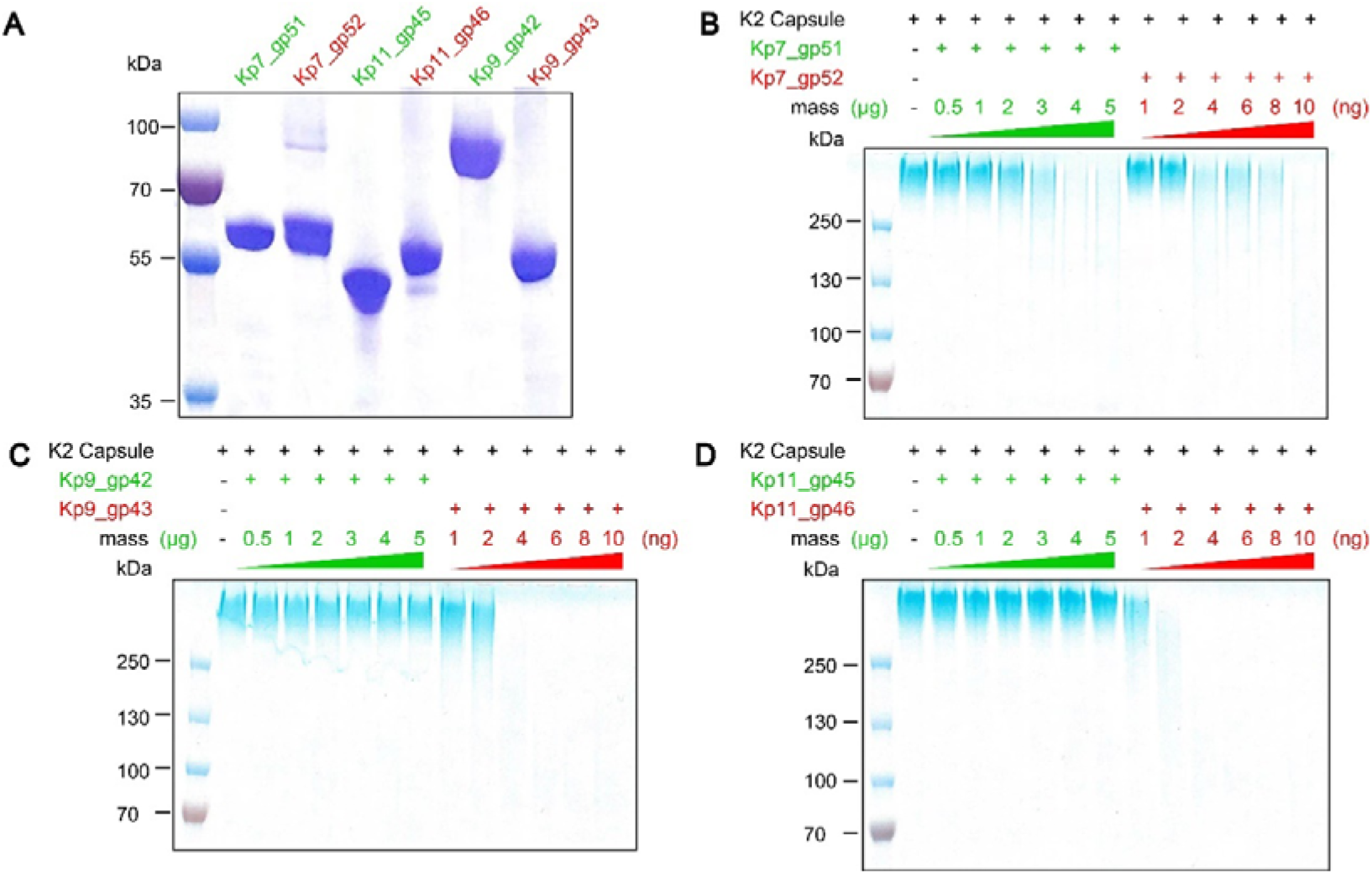
Enzyme activity of tail fibers from Kp7, Kp9, and Kp11. (A) SDS-PAGE gel analysis of the purified phage tail fiber proteins. (B) An Alcian blue stained SDS-PAGE gel showing the purified K2 capsule degraded by tail fiber protein gp51 or gp52 of Kp7. (C) An Alcian blue stained SDS-PAGE gel showing the K2 capsule after being treated with gp42 or gp43 of Kp9. (D) An Alcian blue stained SDS-PAGE showing the purified K2 capsule after being treated with gp45 or gp46 of Kp11.

### 3.7. Mechanisms involved in phage resistance

We observed different susceptibility to phage infection among different clinical isolates with the same K2 serotype capsule. To rule out possible subtle differences among the K2 capsules from the sensitive and insensitive strains, we extracted the K2 capsule and tested the sensitivity of the capsules to the tail fiber protein. The results showed that all the extracted capsules can be degraded by the tail fiber proteins of Kp7 with no evident difference (Supplementary Figure S5). In addition, plaque assays showed that the adsorption process was unaffected in sensitive and insensitive strains (Supplementary Figure S6). To further investigate possible intrinsic mechanisms, we selected representative sensitive and insensitive *K. pneumoniae* isolates and extracted total RNA at different time points post infection (Figure 7A). The extracted total RNA was quantified for bacterial and phage genome by qRT-PCR with corresponding primers (Table S4). The results showed that mRNA levels of the phage encoded essential early genes *orc* and *RNA polymerase* remain unchanged even 15 mins post infection in insensitive strains TH12887 TH13026, TH13030, TH13044 and TH13148 (Figure 7B). whereas mRNA levels of *orc* and *RNA polymerase* have an exponential increase in sensitive strains TH13157 and TH15628. These data combined indicate a transcriptional inhibition of phage encoding essential genes at the early stage of infection in insensitive strains.

**Figure 7.**
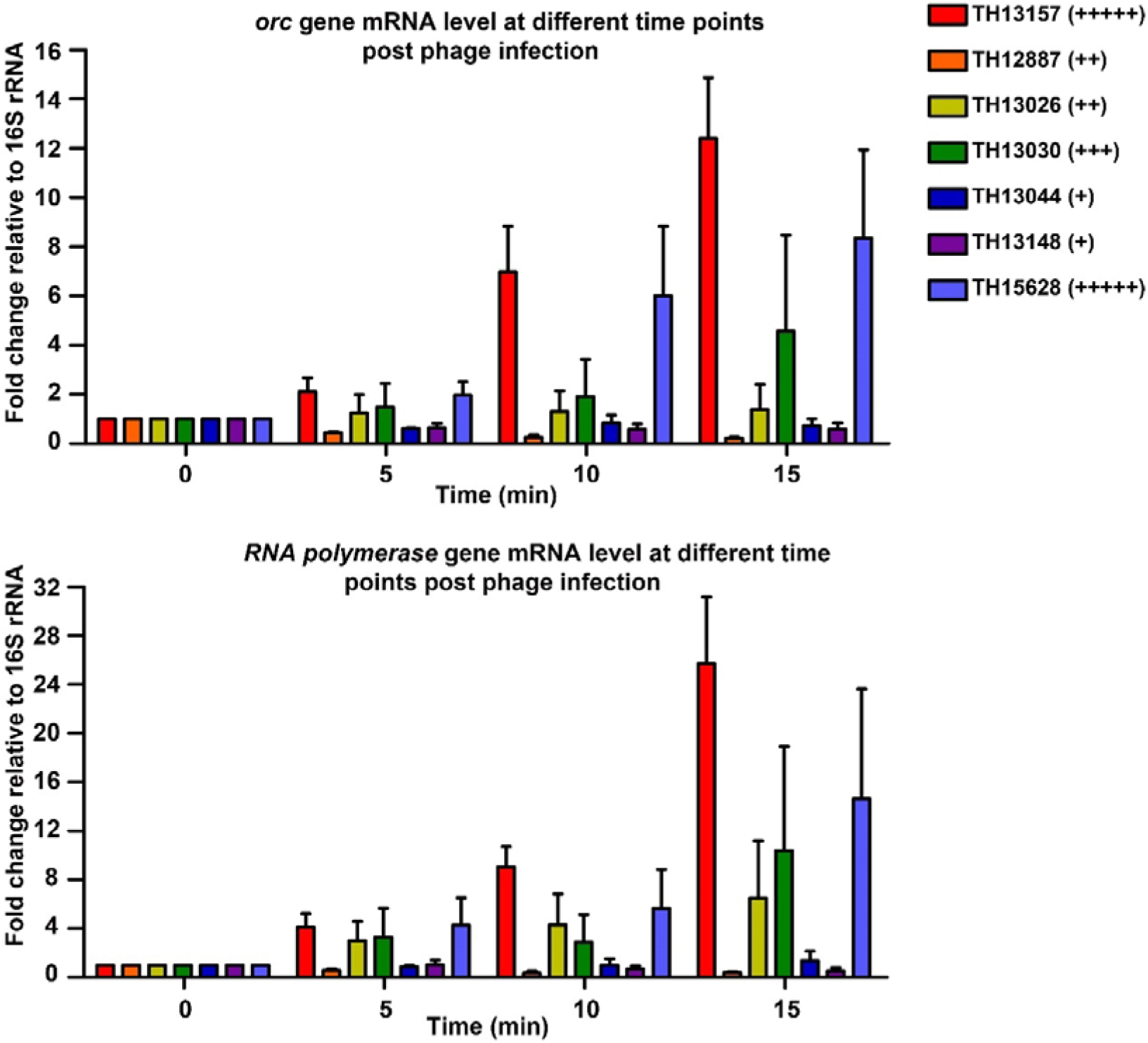
Transcriptional inhibition of key phage genes in insensitive strains. Diagrams showing post infection mRNA levels of phage genes *orc* and *RNA polymerase* in sensitive strains (TH13157, TH15628) and insensitive strain (TH12887, TH13026, TH13030, TH13044, TH13148). The data were normalized against the value at 0 min and were presented as relative fold changes.

### 3.8. Statistical analysis

All the statistical results were analyzed with GraphPad Prism software using Student’s t-test. Differences were considered statistically significant at P< 0.05.

## 4. Discussion

In this study, we isolated, identified, and characterized three bacteriophages that infect and lysate multidrug-resistant *K. pneumoniae* with the K2 capsule. *K. pneumoniae* accounts for a large number of hospital-acquired infections every year. *K. pneumoniae* is featured with the capacity of quickly obtaining drug resistant genes and highly diverse capsule coated on the surface, which is closely related to the adaptation and virulence of this bacterial pathogen. In this study, we confirmed the essential role of *K. pneumoniae* capsules in phage infection. Phage tail fiber proteins that specifically recognize and degrade polysaccharides of *K. pneumoniae* capsules could be used as potential probes for the classifications of *K. pneumoniae*.

Most T7-like podophages have only one tail fiber protein for host recognition. Some podophages, such as SP6 and CBA120, have more than one type of tail fiber proteins assembled for the infection of different host cell strains. For bacteriophages with multiple types of tail fiber proteins, they could have a better chance for dealing with the variations of the host cells and broadening its host tropism. In this study, three lytic bacteriophages (Kp7, Kp9, and Kp11) we identified all have two different types of tail fibers. We proved that both tail fibers of Kp7 could participate in the host bacteria K2 capsule degradation, which indicates that two tail fibers may act synergistically in infection and host cell recognition. However, the tail fiber proteins gp42 of Kp9 and gp45 of Kp11 didn’t show any enzymatic activity towards K2 capsule, suggesting that they may be responsible for recognition of other unidentified capsules.

Phage resistance is a common phenomenon during phage infection. Phage resistance can easily arise due to the receptor mutations, or other systems targeting the different processes of life cycle^20^. In this study, we observed transcriptional inhibition at the early stage of phage infection in insensitive *K. pneumoniae*. Despite the specific interplay between phage and host is unclear, the observation provided insights into possible non-canonical mechanisms of resistance among the clinical isolates of *K. pneumoniae*.

## Author Contributions

Y.X. and J.R.Z. designed the research; L.H., X.T.H. and T.F.Z. performed the experiments; L.H., J.R.Z. and Y.X. analyzed the data and wrote the paper. All authors contributed to the editing of the manuscript.

## Institutional Review Board Statement

Not Applicable.

## Informed Consent Statement

Not Applicable.

## Data availability statement

Whole-genome sequences of *K. pneumoniae* phage Kp7, Kp9, and Kp11 have been deposited in NCBI GenBank under accession numbers ON148527, ON148529, ON148528. All relevant data are provided within the paper and its supplemental material.

## Acknowledgements

We thank Zhiqiang Zhao for his support. We thank the Tsinghua University Branch of the China National Center for Protein Sciences (Beijing) for providing the facility support.

---

**Figure S1.**
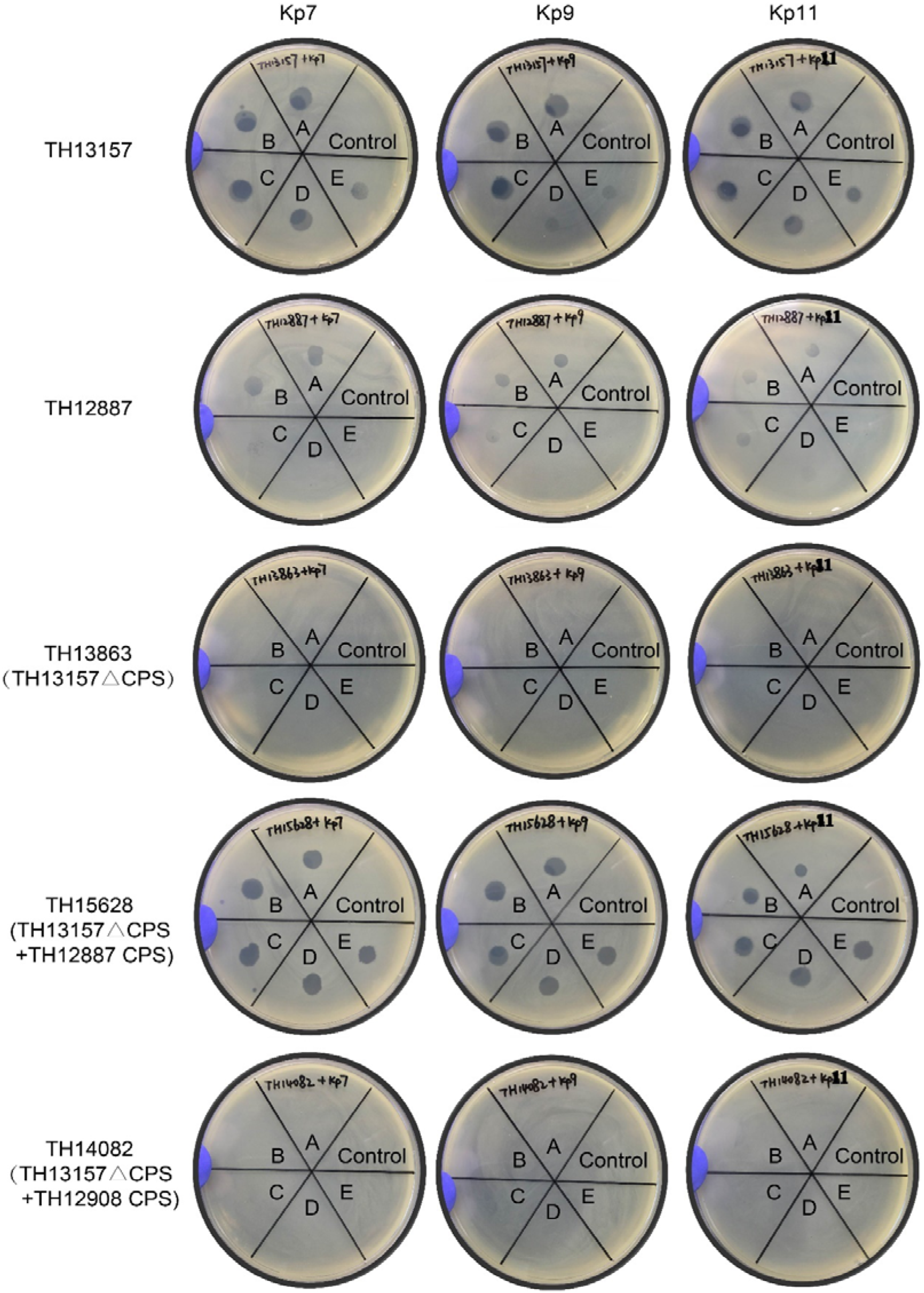
Susceptibility of genetically modified *K. pneumoniae* strains to phage infection. Serial dilutions of Kp7, Kp9, and Kp11 phage (A: 10^10^ pfu/mL, B: 10^9^ pfu/mL, C: 10^8^ pfu/mL, D: 10^7^ pfu/mL, E: 10^6^ pfu/mL) were used for spot tests to identify the phage susceptibility of *K. pneumoniae* strains with genetically modified capsules.

**Figure S2.** Sequence alignments of tail fiber proteins that have enzymatic activity in degrading the isolated K2 capsule. The protein sequences used are Kp7-gp52, Kp9-gp43, Kp11-gp46 and CBA120-orf213. Completely conserved residues are shown in white on a red background. Conserved residues are boxed.

**Figure S3.** Sequence comparisons of tail fiber proteins Kp9-gp42, Kp11-gp45 and KP32-gp37. Completely conserved residues are shown in white on a red background. Conserved residues are boxed.

**Figure S4.**
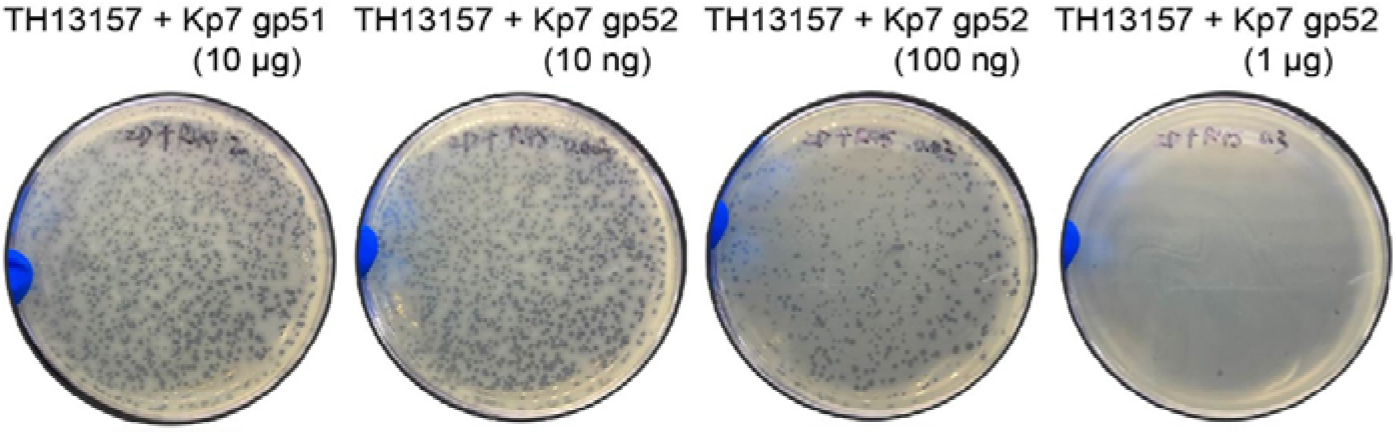
Effects of Kp7-gp51 and Kp7-gp52 on Kp7 phage infection. Plaque assays were performed with *K. pneumoniae* pre-incubated with corresponding Kp7 tail fiber proteins at indicated amount for 1 hour at 37 □.

**Figure S5.**
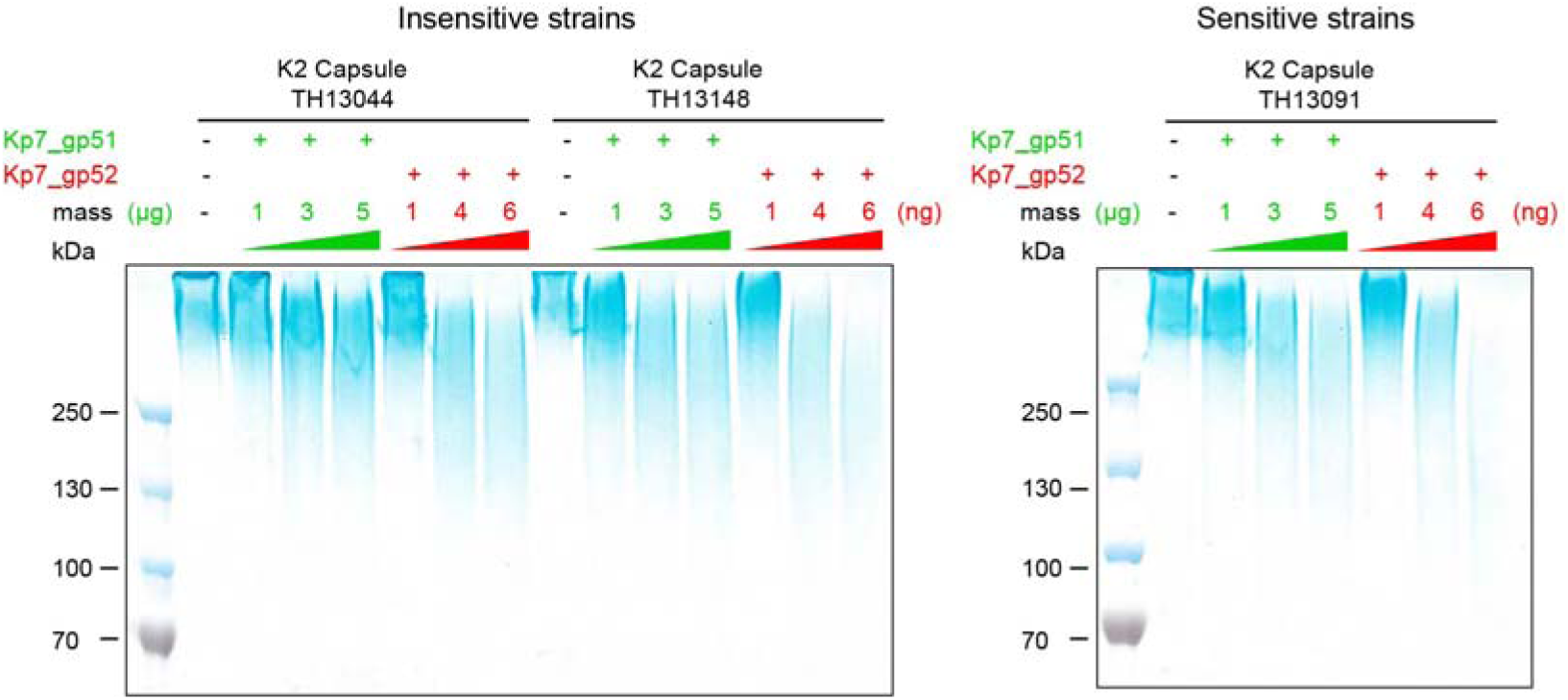
Comparisons of capsular polysaccharides from sensitive (TH13091) and insensitive strains (TH13044 and TH13148). Purified capsular polysaccharides were co-incubated with serial diluted Kp7 tail fiber proteins for 1 hour at 37 □. The mixtures were then analyzed by SDS-PAGE gels stained with Alcian blue.

**Figure S6.**
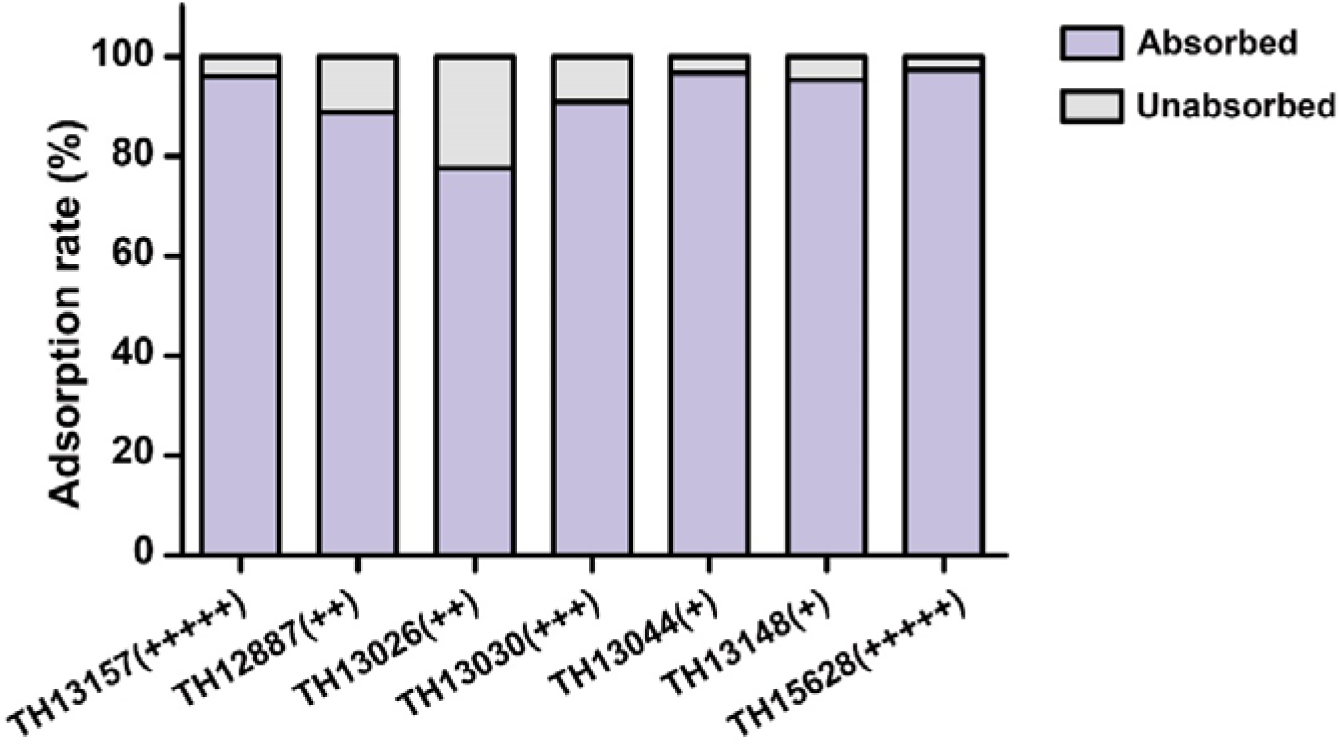
Phage titers in supernatants showing the adsorption efficiency of different *K. pneumoniae* isolates. After adsorption, *K. pneumoniae* cells and phages attached on the cells were pelleted by centrifugation, phage titers in the supernatants were determined by plaque assays. Values in all the assays are the mean values of three measurements.

**Table S1.** Host range of the isolated phages Kp7, Kp9, and Kp11.

| OD<sub>600</sub> | Strain | K<br>type | Susceptibility<br>to phage |  |  | OD<sub>600</sub> | Strain | K<br>type | Susceptibility<br>to phage |  |  |
| --- | --- | --- | --- | --- | --- | --- | --- | --- | --- | --- | --- |
|  |  |  | Kp7 | Kp9 | Kp11 |  |  |  | Kp7 | Kp9 | Kp11 |
| 0.748 | TH12837 | K1 | R | R | R | 0.722 | TH12914 | — | R | R | R |
| 0.713 | TH12849 | K3 | R | R | R | 0.785 | TH12916 | — | R | R | R |
| 0.711 | TH12850 | K47 | R | R | R | 0.732 | TH12973 | K15<br>K17 | R | R | R |
| 0.766 | TH12851 | K47 | R | R | R | 0.785 | TH12974 | K3 | R | R | R |
| 0.784 | TH12852 | K23 | R | R | R | 0.741 | TH12975 | K47 | R | R | R |
| 0.790 | TH12853 | K1 | R | R | R | 0.729 | TH13004 | — | R | R | R |
| 0.742 | TH12881 | K57 | R | R | R | 0.784 | TH13006 | — | R | R | R |
| 0.782 | TH12886 | — | R | R | R | 0.715 | TH13008 | K1 | R | R | R |
| 0.719 | TH12889 | — | R | R | R | 0.779 | TH13012 | K21 | R | R | R |
| 0.756 | TH12890 | K14<br>K64 | R | R | R | 0.716 | TH13013 | — | R | R | R |
| 0.718 | TH12894 | K47 | R | R | R | 0.773 | TH13014 | — | R | R | R |
| 0.735 | TH12895 | K47 | R | R | R | 0.739 | TH13015 | — | R | R | R |
| 0.710 | TH12897 | K47 | R | R | R | 0.786 | TH13016 | K2 | S | S | S |
| 0.743 | TH12898 | — | R | R | R | 0.750 | TH13017 | — | R | R | R |
| 0.717 | TH12899 | — | R | R | R | 0.724 | TH13023 | — | R | R | R |
| 0.711 | TH12900 | — | R | R | R | 0.718 | TH13025 | — | R | R | R |
| 0.729 | TH12904 | — | R | R | R | 0.732 | TH13028 | — | R | R | R |
| 0.790 | TH12905 | — | R | R | R | 0.771 | TH13031 | — | R | R | R |
| 0.744 | TH12909 | — | R | R | R | 0.713 | TH13032 | — | R | R | R |
| 0.712 | TH12912 | — | R | R | R | 0.736 | TH13033 | — | R | R | R |
| 0.758 | TH12913 | — | R | R | R | 0.724 | TH13044 | K2 | S | S | S |

※ R: indicating resistant to phage infection
S: indicating sensitive to phage infection
— : indicating undetermined

**Table S2.**
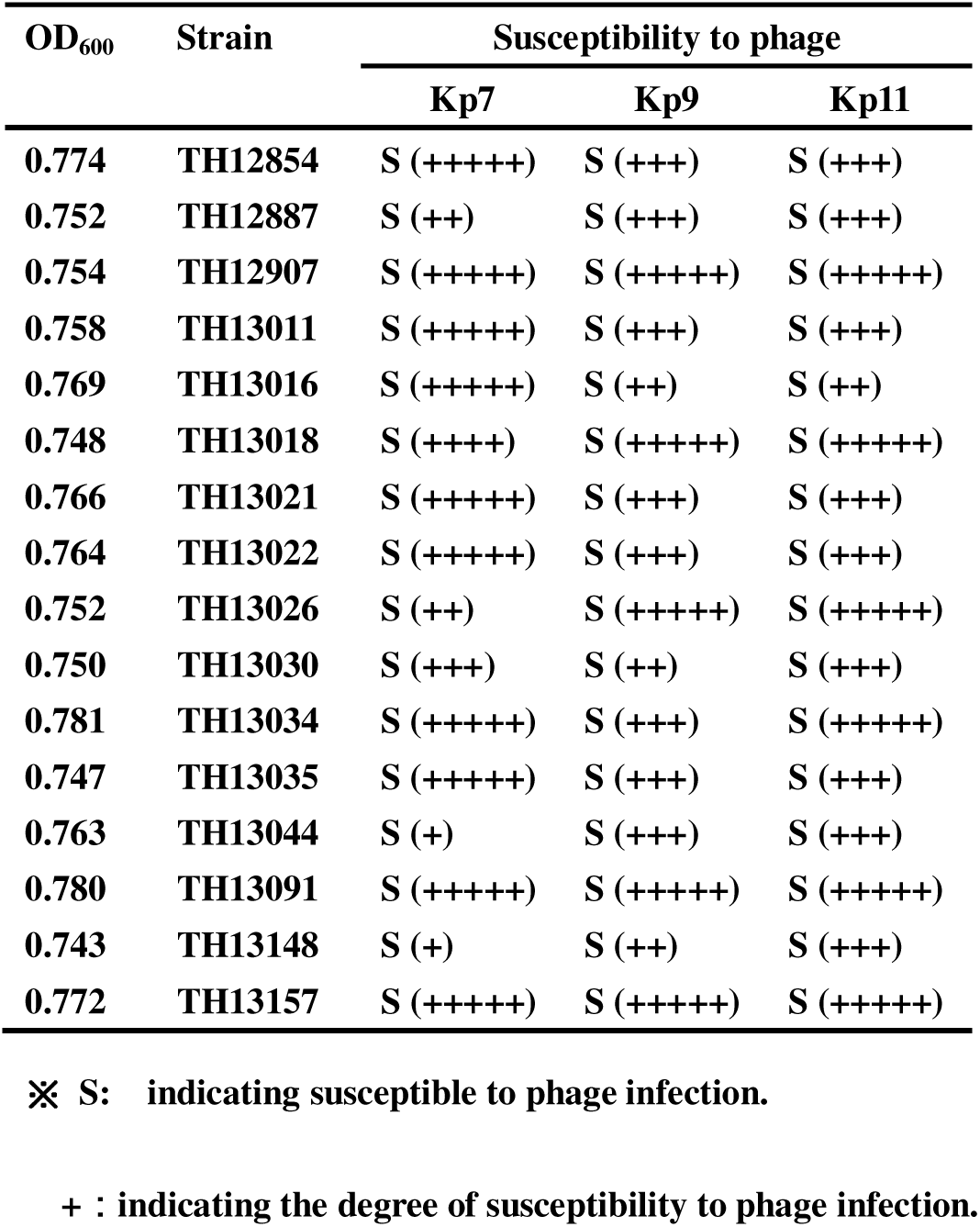
Phage susceptibility towards different isolates of *K. pneumoniae* with K2 capsule.

**Table S3.**
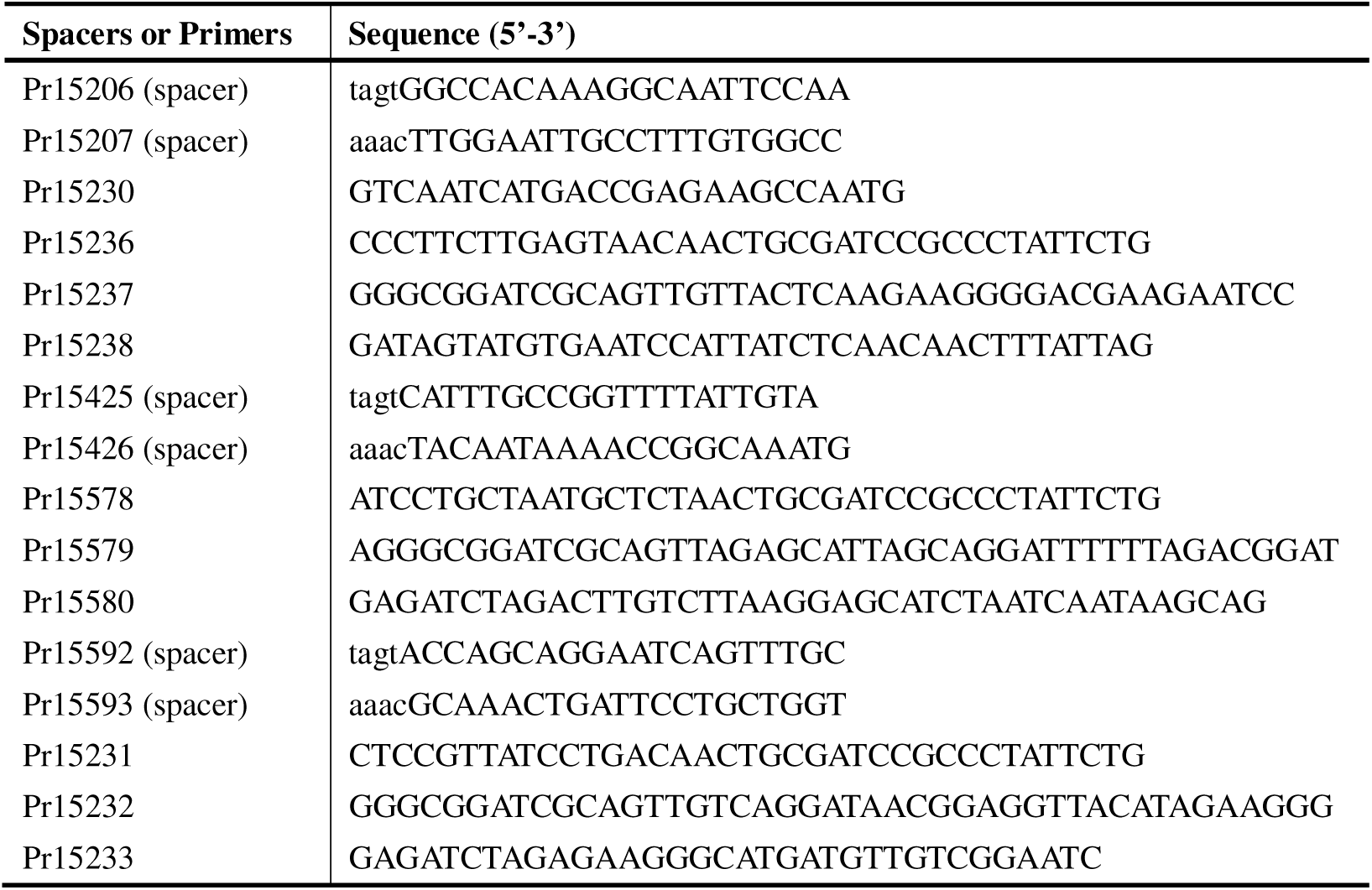
Spacers and primers for constructing strain TH13863.

**Table S4.**
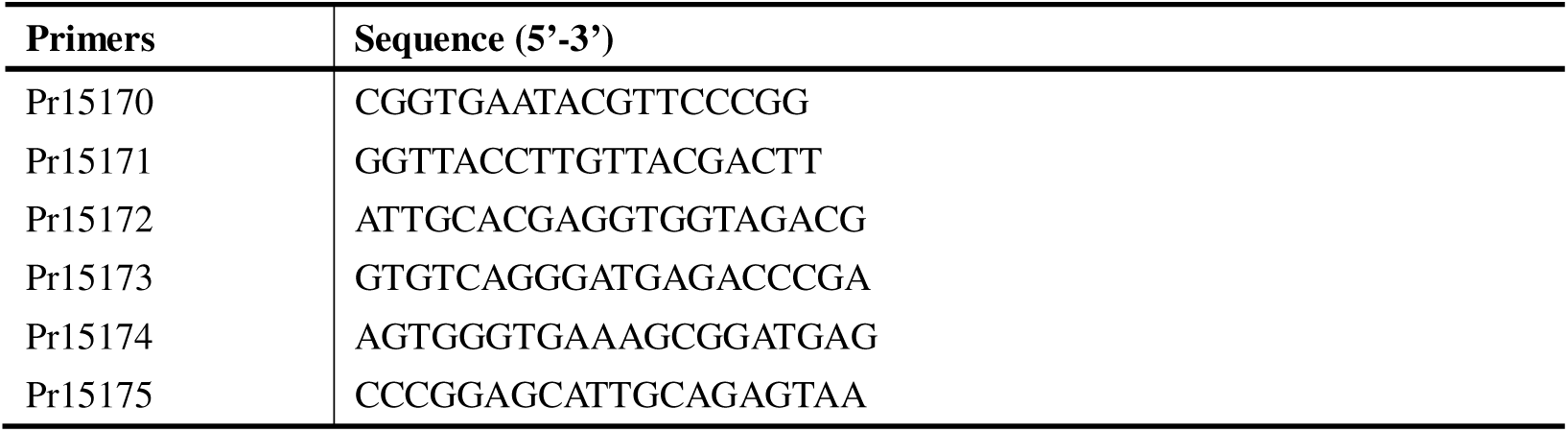
Primers for qRT-PCR.

**Table S5.** Key genomic features of Kp7, Kp9, and Kp11.

| **Phage** | **Genome**<br>**Length (bps)** | **G+C**<br>**Content (%)** | **Number of**<br>**Predicted ORFs** | **Terminal**<br>**Repeat Number** | **Terminal Repeat**<br>**Length (bps)** |
| --- | --- | --- | --- | --- | --- |
| **Kp7** | **45997** | **51.9** | **53** | **2** | **238** |
| **Kp9** | **40337** | **53.1** | **47** | **2** | **177** |
| **Kp11** | **39935** | **53.1** | **50** | **2** | **179** |

